# *Micro*-pellet culture reveals that bone marrow mesenchymal stromal cell (BMSC) chondrogenic induction is triggered by a single day of TGF-β1 exposure

**DOI:** 10.1101/853556

**Authors:** Kathryn Futrega, Pamela G. Robey, Travis J. Klein, Ross W. Crawford, Michael R. Doran

## Abstract

Despite immense promise, engineering of stable cartilage tissue from bone marrow-derived stromal cells (BMSCs, also known as bone marrow-derived “mesenchymal stem cells”) remains elusive. Relative cartilage-like matrix deposition is commonly used to guide BMSC chondrogenic optimisation efforts. However, matrix deposition is heterogeneous in most models, and notably, it lags behind cell fate decisions. We reason that the lag time between cell fate decision and matrix accumulation, coupled with matrix heterogeneity, has obscured basic BMSC biological characteristics, such as differentiation kinetics. Here, we utilize a customized microwell platform to assemble hundreds of small-diameter BMSC *micro*-pellets and characterized chondrogenic differentiation kinetics in response to the canonical signaling molecule, transforming growth factor-β1 (TGF-β1). *Micro*-pellets provide a homogeneous readout, and our experimental design accounts for the significant time delay between growth factor signal and deposition of cartilage-like matrix. While 14-to-21-day induction protocols are routine, BMSC *micro*-pellet cultures reveal that a single day of TGF-β1 exposure was sufficient to trigger chondrogenic differentiation cascades resulting in outcomes similar to *micro*-pellets exposed to TGF-β1 for 21 days. RNA-sequencing analysis demonstrated that one day of TGF-β1 exposure was also sufficient to induce hypertrophic cascades in BMSC, not observed in articular chondrocytes. Refocusing chondrogenic induction optimisation efforts from weeks to the first hours or days of culture, using homogeneous model systems, may benefit efforts to build stable cartilage formed by BMSCs.

**Significance:** The *macro*-pellet model, and assumptions generated using it, have permeated BMSC-based cartilage tissue engineering strategies since the 1990s. Using a *micro*-pellet model, we show that BMSC chondrogenic kinetics are significantly more rapid than historical *macro*-pellets data suggests, and that BMSC chondrogenic and hypertrophic commitment is instructed by a single day of TGF-β1 exposure. This highly relevant study demonstrates that: (1) *macro*-pellets, which are large heterogeneous tissue models confound the differentiation kinetics visible in *micro*-pellet models; (2) induction strategies should focus on the first hours or days of culture; (3) even a single day of TGF-β1 exposure drives BMSC to form hypertrophic tissue *in vivo*, requiring early intervention to prevent hypertrophy; and (4) articular chondrocytes and BMSCs respond distinctly to TGF-β1.

## Introduction

Bone marrow-derived stromal cells (BMSCs, also known as bone marrow-derived “mesenchymal stem cells”) were heralded as a panacea for cartilage repair, but have failed to live up to expectations [1]. While BMSCs appear to have the capacity to differentiate into chondrocytes, current protocols yield temporary, unstable cell populations that evolve to form hypertrophic chondrocytes and mineralized tissue *in vivo* [1, 2]. Because cartilage is matrix-rich (98% [3]) and matrix underpins mechanical function, chondrogenic assays are biased toward the use of matrix characterization as a measure of BMSC differentiation. However, matrix accumulation takes time, and thus matrix accumulated at culture endpoint likely reflects cell fate decisions made days earlier. This temporal complexity is frequently exasperated by the use of macroscopic cartilage tissue models that suffer from profound diffusion gradients, yielding both heterogenous matrix and cell phenotypes [4, 5].

Examination of the classic “pellet culture” highlights how temporal and spatial heterogeneity can confound the study of BMSC chondrogenesis. In 1998, Johnstone *et al.* described the pellet culture (hereafter, referred to as the ***macro*-pellet**), which has become the gold standard for differentiating BMSCs into chondrocyte-like cells *in vitro* [6]. In the original *macro*-pellet culture, ~2×10^5^ BMSC were pelleted in induction medium that critically included supplementation with transforming growth factor-β1 (TGF-β1). Thousands of papers have since used the *macro*-pellet culture to characterize or optimise the *in vitro* chondrogenic capacity of BMSCs. Little has changed since the introduction of the *macro*-pellet culture model, with most studies continuing to use similar numbers of cells per *macro*-pellet, and several weeks of induction in medium supplemented with one of the TGF-β isoforms (TGF-β1 or TGF-β3). Recent reviews of published chondrogenic induction models described the use of TGF-β induction ranging from 7 days to 28 days, with *macro*-pellets containing between 2×10^5^ to 1×10^6^ BMSCs each [7, 8]. Other growth factors have been incorporated to enhance chondrogenic induction protocols [7–11], mostly in *macro*-pellet cultures formed from >2×10^5^ BMSCs each and induction periods ranging from 7-45 days; however, TGF-β remains the most important and most commonly used chondrogenic induction factor.

TGF-β instructs BMSCs to take on a chondrocyte-like phenotype, and in response, pelleted cells secrete cartilage-like matrix. *Macro*-pellet cultures evolve to form large (1-3 mm diameter) tissues, yielding large radial diffusion gradients and the development of different cell and tissue types in different regions of the pellet [4, 5]. The radially heterogeneous matrix distribution that can be observed at different time points suggest that some regions of the *macro*-pellet undergo differentiation at different rates than other regions of the *macro*-pellet [5]. While spatial heterogeneity is well-recognized, the potential impact that this imposes on temporal differentiation heterogeneity is rarely discussed. As BMSC chondrogenic differentiation lies on a continuum, where the terminally differentiated cell type is a hypertrophic chondrocyte [12], the temporal response to induction factors is a critical variable. Geometric heterogeneity common in *macro*-pellet cultures, and other macroscopic tissue models, effectively imposes temporal heterogeneity. Because of these uncontrolled gradients, we reason that fundamental understanding, such as true BMSC induction kinetics in response to factors such as TGF-β1 remains unknown. Lack of understanding of induction kinetics has hindered development of protocols that might better control stage-wise differentiation of BMSCs into a cell population capable of producing hyaline cartilage, with reduced propensity to undergo hypertrophic differentiation. Effective bioprocess optimization will require use of platforms that enable manufacture of more homogeneous cartilage-like tissues to quantify induction kinetics and more precisely guide desired tissue outcomes.

Here, we use a more homogeneous, small diameter *micro*-pellet model to characterize the temporal influence of TGF-β1, the most commonly used BMSC chondrogenic induction factor. We used a high-throughput microwell platform, termed the *Microwell-mesh* [5], to enable manufacture of thousands of small diameter *micro*-pellets for this analysis. Because individual *micro*-pellets (5×10^3^ cells each) are formed from fewer cells than *macro*-pellets (2×10^5^ cells each), the resulting smaller tissues experience reduced diffusion gradients and yield more homogeneous cartilage-like tissue. For this reason, *micro*-pellets are a more appropriate model to characterize BMSC chondrogenic differentiation kinetics, compared with the standard *macro*-pellet model. To account for the disconnect or lag time between cellular response to TGF-β1 exposure and cartilage-like matrix accumulation, we used 21-day cultures, where cells were exposed to TGF-β1 for different time periods ranging from 0, 1, 3, 7, 14, or 21 days. We contrasted results from *micro*-pellet and *macro*-pellet tissues formed from either BMSCs or *in vitro* expanded articular chondrocytes (ACh). We performed RNA sequencing of *micro*-pellet cultures at various days (0, 1, 3, 7, and 21) of the 21-day culture period and on day 21 following TGF-β1 withdrawal (on day 1, 3, and 7). These data reveal the time-dependent cellular response to TGF-β1 signalling, as well as provide insight into how different durations of TGF-β1 exposure influences BMSC fate. Overall, *micro*-pellet studies reveal the rapid temporal response of BMSCs to TGF-β1 exposure, the influence of TGF-β1 programming following TGF-β1 withdrawal, the confounding influence that *macro*-pellet heterogeneity has on perceived differentiation kinetics, and the divergent response of ACh and BMSCs to the TGF-β1 stimulus.

## Results

TGF-β1 is the most common growth factor used in BMSC chondrogenic induction medium. However, because of confounding factors in chondrogenic models, the temporal response of BMSCs to TGF-β1 and BMSC differentiation kinetics are uncertain. To determine the temporal influence of TGF-β1 on BMSC chondrogenic induction, we tested various TGF-β1 exposure times over a fixed culture period of 21 days to account for the lag between cell differentiation and measurable matrix accumulation. Traditional *macro*-pellet cultures were assembled from 2×10^5^ BMSCs each in deep well plates, while *micro*-pellets were formed from 5×10^3^ BMSCs each (40-fold fewer cells per tissue) in Microwell-mesh plates (**Fig 1A**). In Microwell-mesh plates (**Fig 1B**), cells in suspension were centrifuged through the openings (36 μm) in the nylon meshes bonded over microwells. Cells pelleted in microwells aggregated to form *micro*-pellets, becoming too large to escape back through the mesh openings, thus being retained in discrete microwells over the culture period. By varying the number of cells per pellet, large *macro*-pellets (2×10^5^ BMSC each) or smaller *micro*-pellets (5×10^3^ BMSC each) were produced. Larger diameter pellets inherently suffer from increased diffusion gradients of metabolites, gases, and other factors (**Fig 1C**) [13, 14], while gradients are reduced in smaller diameter pellets, yielding more homogeneous cartilage-like tissue [4, 5]. Induction cultures consisted of basal chondrogenic medium supplemented with TGF-β1 for either 0, 1, 3, 7, 14, or 21 days of the total 21-day induction culture. Each culture condition is represented by a horizontal line in **Fig 1D**, with blue and grey lines representing days with (+)TGF-β1 and without (−)TGF-β1, respectively, and the red arrow specifying the day of analysis. On the indicated day, TGF-β1 was eliminated (washed away) and replaced with basal medium for the remainder of the 21-day culture. Parallel cultures were established using BMSCs from 4 unique donors and ACh from two unique donors.

**Figure 1.**
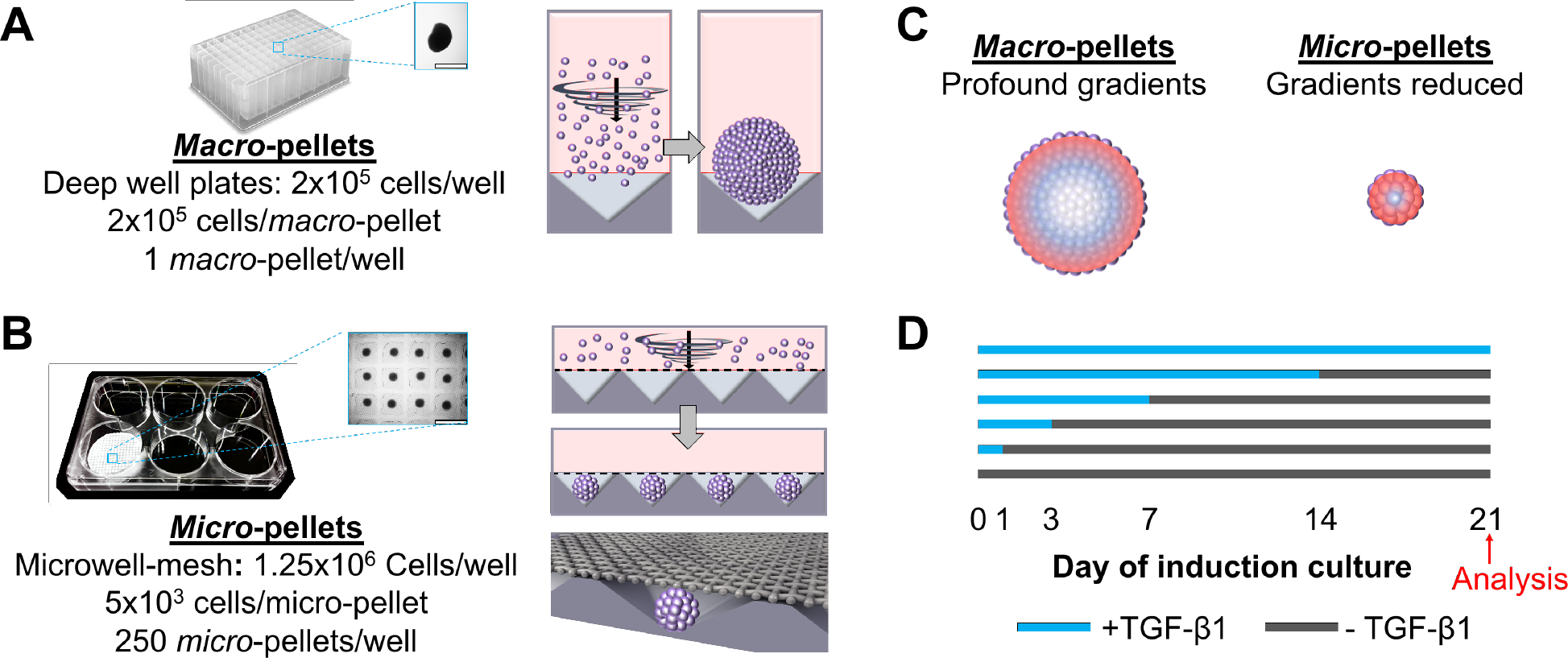
Schematic of experimental procedures. (A, B) Different cell seeding densities were used to generate *macro*-pellets or *micro*-pellets of specific size in deep well plates or Microwell-mesh plates, respectively. *Micro*-pellets were retained in discrete microwells by the nylon mesh bonded over the microwell openings. Retention by the mesh enabled long-term *micro*-pellet culture, including multiple medium exchanges to add or deplete cultures of TGF-β1. (C) Diffusion gradients are reduced in smaller diameter *micro*-pellets relative to larger diameter *macro*-pellets [13, 14]. (D) Cultures were carried out for 21 days total, with blue lines representing culture days with TGF-β1 in the medium, and grey lines representing culture days without TGF-β1 in the medium.

### Gross Morphology and Histology

Gross photographic and microscopic images, and Toluidine blue stained sections for BMSC- and ACh-induced chondrogenic tissues exposed to TGF-β1 for 0, 1, 3, 7, 14, or 21 days are shown in **Fig 2** and **3**, respectively. Overall, tissue growth was greater in *micro*-pellets than in *macro*-pellets (**Fig 2A**). On average, the diameter of *macro*-pellets exposed to TGF-β1 for 21 days was 2.2 times greater than *macro*-pellets not exposed to TGF-β1 (Day 0), while the diameter of *micro*-pellets exposed to TGF-β1 for 21 days was 3.7 times greater in diameter than *micro*-pellets not exposed to TGF-β1 (Day 0). Relative cartilage-like tissue volume was approximately 25% greater per input cell when BMSCs were cultured as *micro*-pellets with 21 days TGF-β1 exposure, relative to matched BMSC *macro*-pellet controls. In *macro*-pellet cultures, tissues grew incrementally larger with extended TGF-β1 exposure. By contrast, *micro*-pellets achieved the majority of tissue growth in response to a single day of TGF-β1 exposure (approximately 75% of maximum *micro*-pellet diameter was achieved in response to a single day of TGF-β1 exposure).

**Figure 2.**
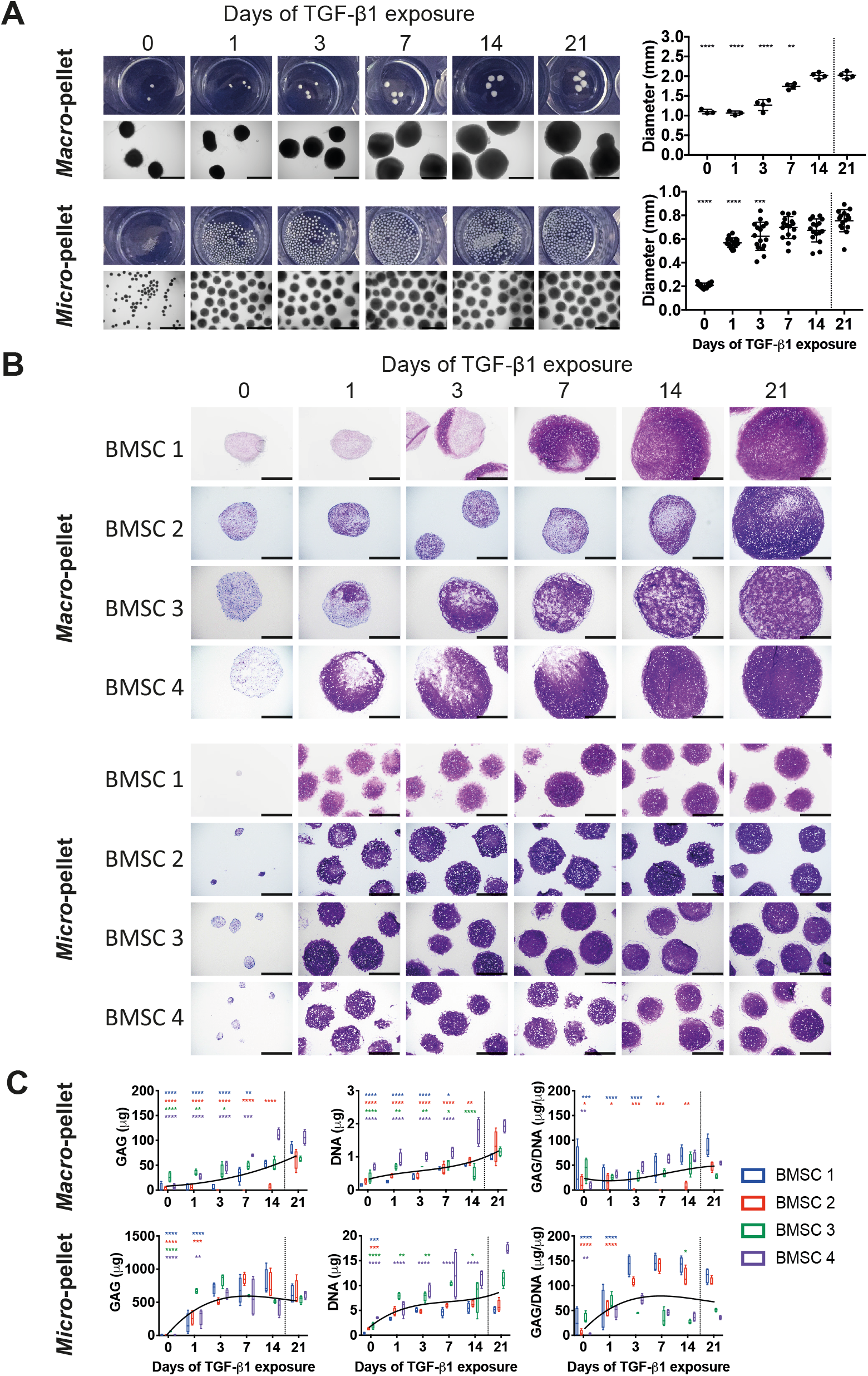
Cartilage-like tissues derived from BMSCs after 21 days *in vitro* differentiation with TGF-β1 exposure for the number of days indicated. (A) Gross photographs and brightfield microscopy images of whole BMSC-derived tissues (scale bars for microscopy images, 1 mm), and corresponding diameters. (B) Toluidine blue stain of *macro*- and *micro*-pellet tissue cross-sections are shown for four unique BMSC donors (scale bars, 500 μm). (C) GAG, DNA, and normalized GAG/DNA quantification for each BMSC donors. Asterisks are shown for values that are significantly lower than the corresponding 21 days of TGF-β1 exposure control. Values represent mean ± SD; n=4; *P < 0.05, **P < 0.01, ***P < 0.001, and ****P < 0.0001.

Toluidine blue histological staining was used to characterize spatial glycosaminoglycan (GAG) content in the tissue matrix. Both BMSC *macro*-pellet and *micro*-pellet cultures that were not supplemented with TGF-β1 during the induction period remained small (**Fig 2A**) and had little GAG staining in their matrix (**Fig 2B**). BMSC *macro*-pellets exposed to TGF-β1 for less than 21 days, simultaneously had regions of undifferentiated tissue (weak Toluidine blue GAG staining) and regions of differentiated tissue (strong Toluidine blue GAG staining) (**Fig 2B**). In BMSC *micro*-pellets, GAG staining was relatively uniform across the diameter of tissues, and strong GAG staining was visible even with a single day of TGF-β1 exposure (**Fig 2B**). Substantial BMSC donor variability was observed in *macro*-pellets when the duration of TGF-β1 exposure was short (i.e. less than 14 days). However, donor variability was less evident in *micro*-pellet cultures, with all BMSC populations yielding tissue that stained intensely and uniformly with Toluidine blue, even with only 1 day of TGF-β1 exposure (**Fig 2B**).

The biochemical content, including GAG, DNA and GAG/DNA, for BMSC *macro*- and *micro*-pellets is shown in **Fig 2C**. Cultures that received TGF-β1 for the full 21 days were considered the control (shown to the right of dashed lines in **Fig 2C**), and cultures with different durations of TGF-β1 exposure were compared with these controls. In BMSC *macro*-pellet cultures, two of four BMSC donors reached maximal GAG content with 14 days of TGF-β1 exposure, while one donor required the full 21 days of TGF-β1 exposure (**Fig 2C**). In BMSC *micro*-pellet cultures, three of four BMSC donors reached maximum GAG content with 3 days of TGF-β1 exposure, while one donor reached this maximum value with 1 day of TGF-β1 exposure. In BMSC *macro*-pellet cultures, DNA content increased steadily with increasing TGF-β1 exposure days, with two donors reaching maximum DNA content with 14 days of TGF-β1 exposure and two requiring 21 days of TGF-β1 exposure. In BMSC *micro*-pellet cultures, DNA content also increased with increasing TGF-β1 exposure, with two donors reaching maximum DNA content with 1 day of TGF-β1 exposure and two donors reaching maximum DNA content with 3 or 7 days of TGF-β1 exposure.

For BMSC *macro*-pellets, maximum GAG/DNA was variable, with two BMSC donors (BMSC 3 and 4) reaching the maximum GAG/DNA content with 1 day of TGF-β1 exposure, and two donors (BMSC 1 and 2) required 14 and 21 days of TGF-β1 exposure, respectively. By contrast, BMSC *micro*-pellets maximum GAG/DNA was achieved with reduced TGF-β1 exposure. BMSC donors 3 and 4 reached maximum GAG/DNA content with 1 day of TGF-β1 exposure, while BMSC donors 1 and 2 required 3 and 7 days of TGF-β1 exposure, respectively. As the GAG and DNA quantification assay was only performed at day 21 of culture, it is not possible to determine at what rate the GAG or DNA content was produced and whether TGF-β1 might influence the rate of GAG or DNA production differently.

Overall, fewer days of TGF-β1 exposure in *micro*-pellet BMSC cultures were required to obtain GAG quantities equivalent to control culture having 21 days of TGF-β1 exposure, compared to traditional *macro*-pellet BMSC cultures. Spatial distribution of Toluidine blue staining demonstrated uniform and high levels of GAG deposition in *micro*-pellet tissues with as little as 1 day of TGF-β1 exposure, while GAG deposition in *macro*-pellets required 14-21 days TGF-β1 exposure before achieving similar uniformity.

ACh *macro*- and *micro*-pellet cultures (**Fig 3**) had a different pattern of behaviour compared with their equivalent BMSC pellet cultures described above. ACh-derived *macro*-pellets appeared similar in size with varied TGF-β1 exposure following 21 days of culture (**Fig 3A**). Unlike spherical BMSC-derived tissues, ACh *macro*-pellets exhibited a cup-like structure with a concaved centre (see red arrows pointing to side views in **Fig 3A**). This cup-like structure was not observed in ACh *micro*-pellets, which were spherical like BMSC *micro*-pellets, but required more than 7 days of TGF-β1 exposure to grow to the same extent as ACh cultures exposed to TGF-β1 for the full 21-day culture period. Like BMSC *macro*-pellets, ACh *macro*-pellets benefited from longer TGF-β1 exposure, as evidenced by the gradual increase in pellet size (**Fig 3A**) and increased GAG content, with increasing days of TGF-β1 exposure (**Fig 3B-C**). However, unlike BMSC *micro*-pellets, ACh *micro*-pellets required prolonged TGF-β1 exposure for increased growth (**Fig 3A**), and GAG production. ACh *macro*- and *micro*-pellets demonstrated reduced GAG staining (**Fig 3B**) and content (**Fig 3C**) when TGF-β1 was withdrawn early in the 21-day culture period. In both *macro*- and *micro*-pellet culture, ACh donor 2 appeared to be a more potent cartilage-matrix producing cell population, compared with ACh donor 1, as donor 2 cells produced more intense GAG staining (**Fig 3B**) and GAG quantity (**Fig 3C**). Culturing ACh 1 as *micro*-pellets did improve the potency of this donor, possibly due to increasing the cells’ proliferative capacity as seen by the substantial increase in DNA content in *micro*-pellets, compared with the very low rate of DNA content increase in the *macro*-pellets (**Fig 3C**). The differences in growth, GAG and DNA production between ACh and BMSC-derived tissues may be explained by the fact that our ACh were of later passage than BMSC (P6 vs. P3, respectively). Additionally, as ACh are considered to be differentiated cells, the mechanisms of TGF-β1 action on these cells are likely to be substantially different than on multipotent BMSCs undergoing a fate choice and differentiation *in vitro*.

**Figure 3.**
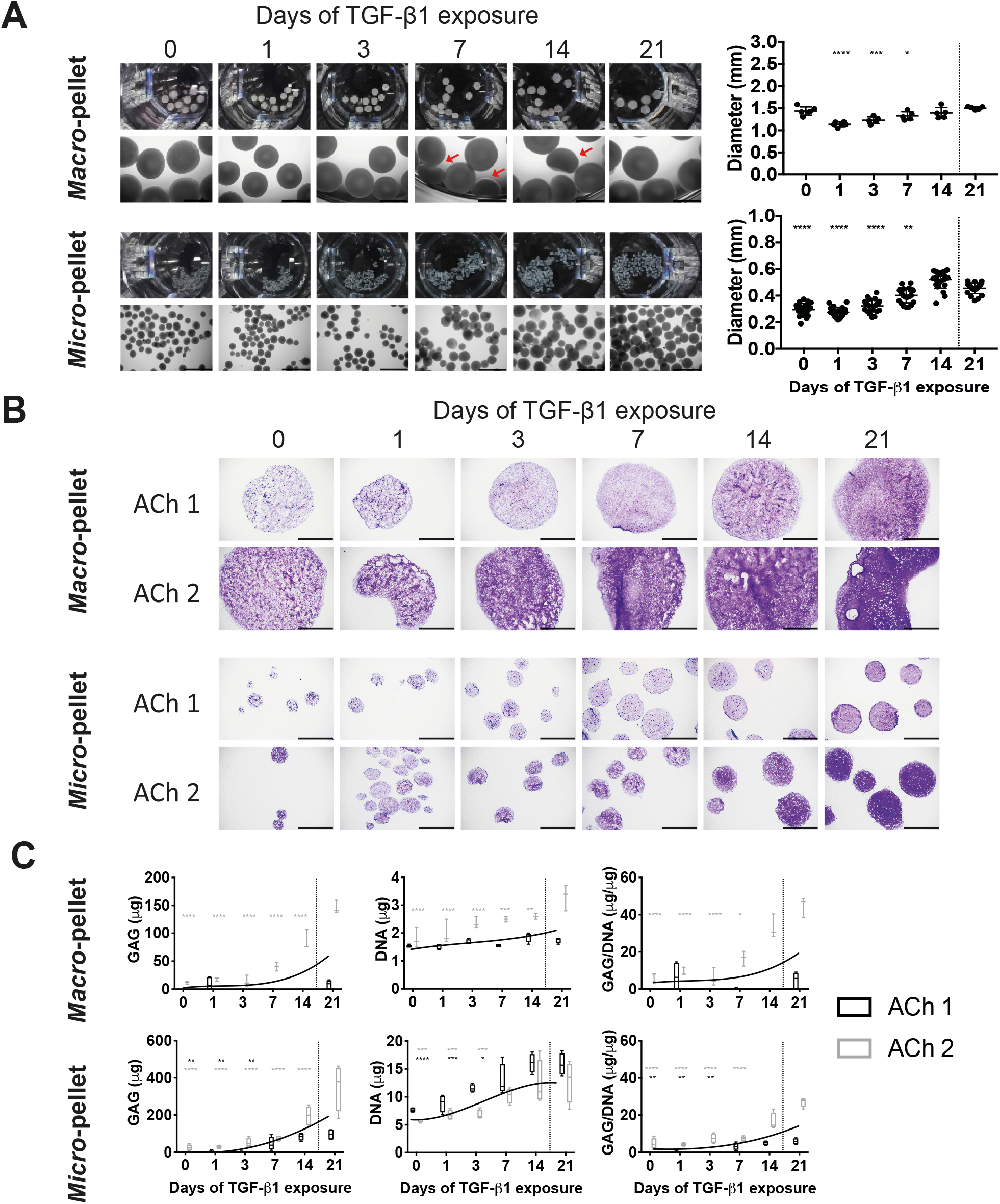
Cartilage-like tissues derived from ACh after 21 days *in vitro* differentiation with TGF-β1 exposure for the number of days indicated. (A) Gross photographs and brightfield microscopy images of whole ACh-derived tissues (scale bars for microscopy images, 1 mm), and corresponding diameters. (B) Toluidine blue stain of *macro*- and *micro*-pellet tissue cross-sections are shown for two unique ACh donors (scale bars, 500 μm). (C) GAG, DNA, and normalized GAG/DNA quantification for ACh donors. Asterisk are shown for values that are significantly lower than the corresponding 21 days of TGF-β1 exposure control. Values represent mean ± SD; n=4; *P < 0.05, **P < 0.01, ***P < 0.001, and ****P < 0.0001.

### qPCR Analysis

We assessed the relative expression of genes associated with chondrogenesis (*SOX9, COL2A1, ACAN*) and hypertrophy (*COL1A1, COL10A1, and RUNX2*) for both *macro*- (**Fig 4A**) and *micro*-pellet (**Fig 4B**) cultures. Each TGF-β1 exposure condition was compared with the standard control culture, where TGF-β1 was maintained for the entire 21 days (shown to the right of dashed lines in **Fig 4**).

**Figure 4.**
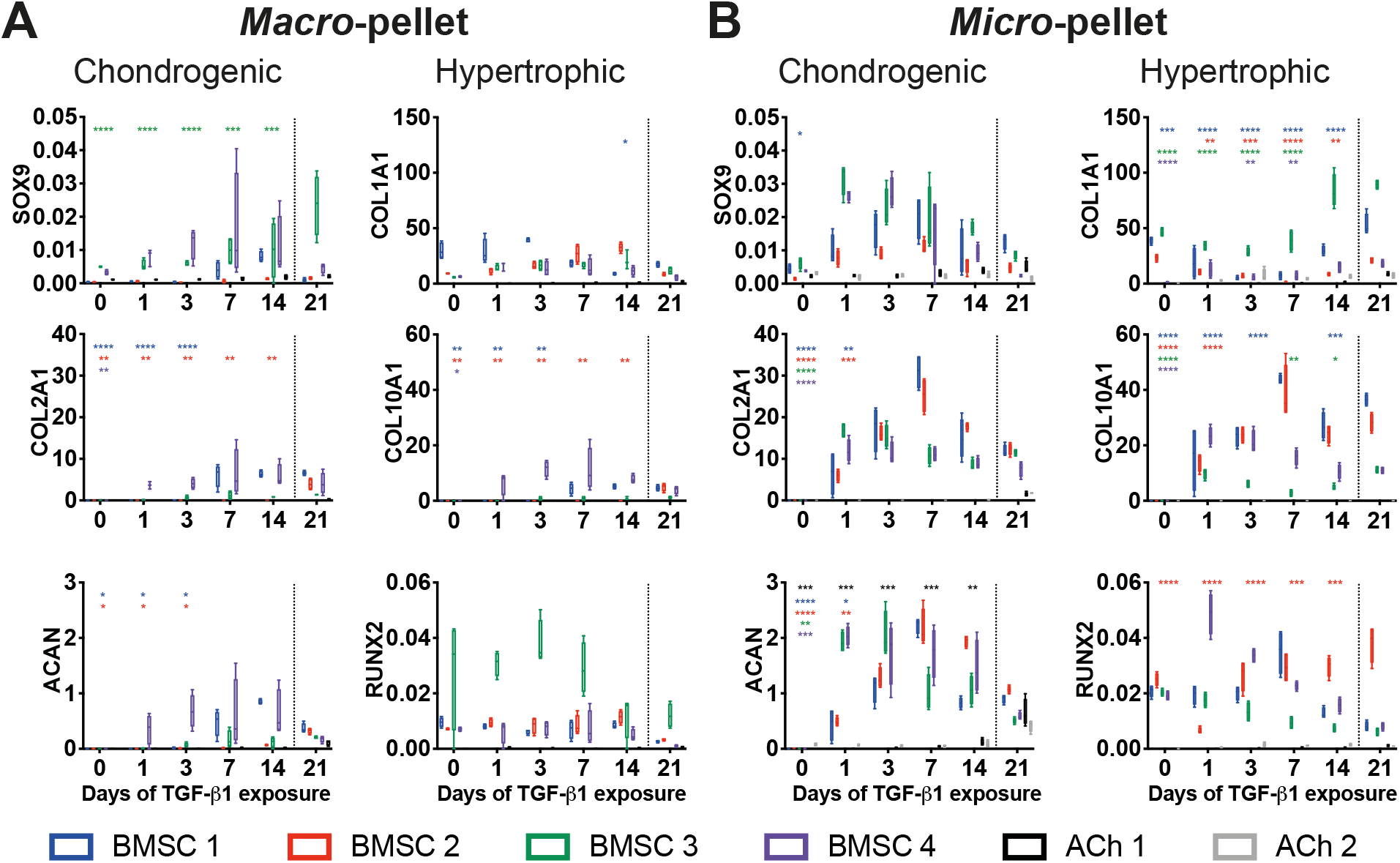
qPCR analysis of cartilage-like tissues derived from BMSCs or ACh on day 21 of culture, following varying days of TGF-β1 exposure. (A) qPCR from *macro*-pellet cultures, (B) qPCR from *micro*-pellet cultures. All values are compared with the standard 21 days of TGF-β1 exposure control (separated by the dotted line). Relative gene expression represents ∆Ct values normalized to GAPDH expression. Asterisks are shown for values that are significantly lower than the corresponding 21 days of TGF-β1 exposure control. Values represent mean ± SD; n=4; *P < 0.05, **P < 0.01, ***P < 0.001, and ****P < 0.0001.

In *macro*-pellets, substantial variability was observed between different cell donors. While both chondrogenic and hypertrophic genes tended to increase with extended TGF-β1 exposure, these were not statistically significant for all donors and at all time points analyzed. *Micro*-pellet cultures yielded more consistent gene expression changes. This outcome can, in part, be related to the greater magnitude in gene expression values observed in *micro*-pellet cultures (**Fig 4**). For each BMSC donor, relative *SOX9* expression was statistically equivalent or greater for cultures that only received 1-14 days of TGF-β1 exposure, compared with the control. Relative *COL2A1* and *ACAN* expression was comparable to the 21-day control condition with 1 day of TGF-β1 exposure for BMSC donors 3 and 4, and 3 days for BMSC donors 1 and 2.

The expression of *COL10A1* in *micro*-pellets formed from BMSC donors 3 and 4 reached 21-day control levels with 1 day of TGF-β1 exposure, while *micro*-pellets formed from BMSC donor 2 required 3 days of TGF-β1 exposure, and BMSC donor 1 required 7 days of TGF-β1 exposure. *RUNX2* expression was only significantly lower for BMSC donor 2 when TGF-β1 was washed out at days 1, 3, 7, or 14, compared with the 21-day control. *COL1A1* expression in BMSC *micro*-pellets was the only gene that declined significantly when TGF-β1 was washed out on one or more of the washout days (day 1, 3, 7, or 14) for all donors. For ACh, *ACAN* expression declined when TGF-β1 was washed out at days 1, 3, 7, or 14, but this was only significant for one of the two ACh donors, ACh 1. No hypertrophy gene expression was observed in cultures from either of the two ACh donors.

Overall, qPCR analysis suggested that 1-3 days of BMSC *micro*-pellet exposure to TGF-β1 was sufficient to induce chondrogenic genes, and this same brief TGF-β1 exposure appeared to also upregulate hypertrophy genes, as measured in the 21-day control cultures.

### *In vivo* subcutaneous hypertrophy

To confirm our qPCR analysis, which showed that even with reduced TGF-β1 exposure, BMSC *micro*-pellets still appeared to express high relative levels of the hypertrophic marker, *COL10A1*, we packed *micro*-pellets into bovine osteochondral defect models and transplanted them subcutaneously in NSG mice for 8 weeks using a previously established method [15]. Histological examination of tissues showed that, unlike ACh-derived *micro*-pellets, BMSC-derived *micro*-pellets exhibited a hypertrophic appearance based on diminished GAG staining and significant recruitment of blood cells, forming a bone marrow like tissue (**Fig 5A**). Micro-CT analysis confirmed that, unlike differentiated ACh *micro*-pellets, BMSC *micro*-pellets formed mineralized tissue *in vivo* regardless of TGF-β1 exposure duration (**Fig 5B**). *Micro*-pellets exposed to TGF-β1 for 3 days yielded mineralized tissue, with a density similar to the bovine subchondral bone contained in the plug of the joint used to develop the artificial cartilage defect model. ACh *micro*-pellets formed continuous fill, but individual *micro*-pellets were still discernible. While not mineralized, the GAG staining of ACh *micro*-pellets was faint, suggesting that the cartilage-like tissue may be low quality. It may be possible to improve the tissue quality by using lower passage ACh; however, maintaining low passage ACh, while generating sufficient cartilage-defect fill material is a significant limitation in the use of ACh.

**Figure 5.**
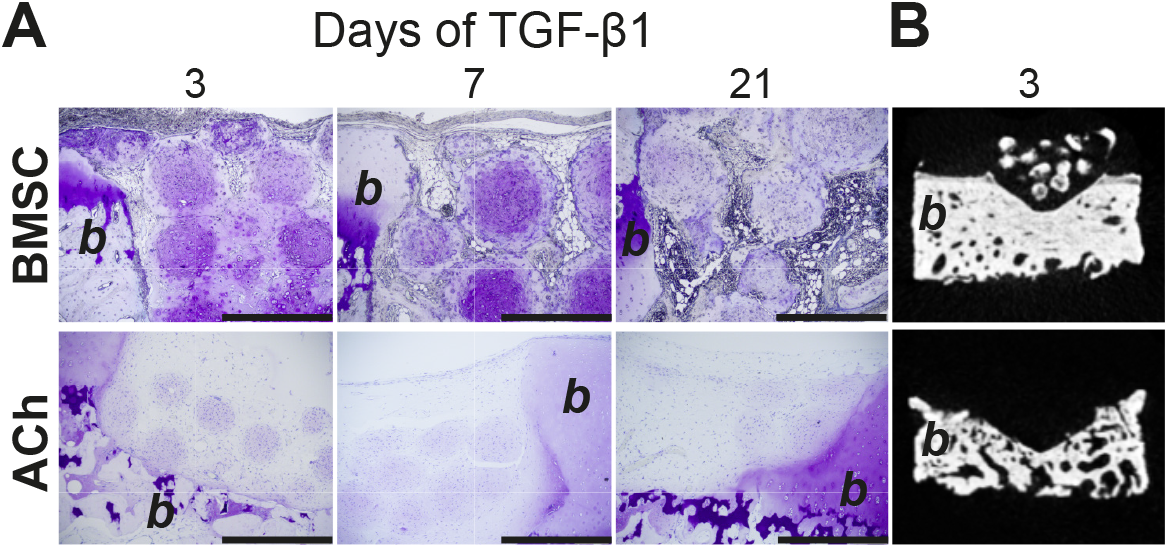
Analysis of *micro*-pellets implanted subcutaneously in NSG mice for 8 weeks in bovine defect models. (A) Histological sections showed that BMSC and ACh *micro*-pellets appeared highly remodelled with reduced GAG staining (Toluidine blue), and only BMSC formed bone marrow-like tissues (scale bars, 500 μm). (B) Micro-CT analysis confirmed the presence of mineralized tissue in BMSC *micro*-pellets, unlike ACh *micro*-pellets. Representative images are shown for 21 day *in vitro* differentiated tissues with TGF-β1 exposure for days indicated on top of the images. Surrounding bovine cartilage and bone tissue is marked with the letter “*b*” to indicate bovine tissue.

### RNA-Seq Analysis

To investigate which genes where modified by TGF-β1 in BMSC and ACh cultures and identify potential targets to inhibit hypertrophy, we performed bulk RNA-seq analysis. We analysed cultures before induction (Day 0) and harvested on day 1, 3, 7, and 21 of culture induction in the presence of TGF-β1. On each day, we analysed three BMSC donors (BMSC 1, 2, and 3) and two ACh donors (ACh 1 and 2). Additionally, we analyzed one BMSC donor (BMSC 1) and one ACh donor (ACh 2) on day 21 that was exposed to TGF-β1 for 1, 3, or 7 days of the total 21-day culture. A schematic of the experiment is shown in **Fig 6A** with blue and grey lines representing culture days with (+)TGF-β1 and without (−)TGF-β1, respectively, and red arrows representing the day of analysis.

**Figure 6.**
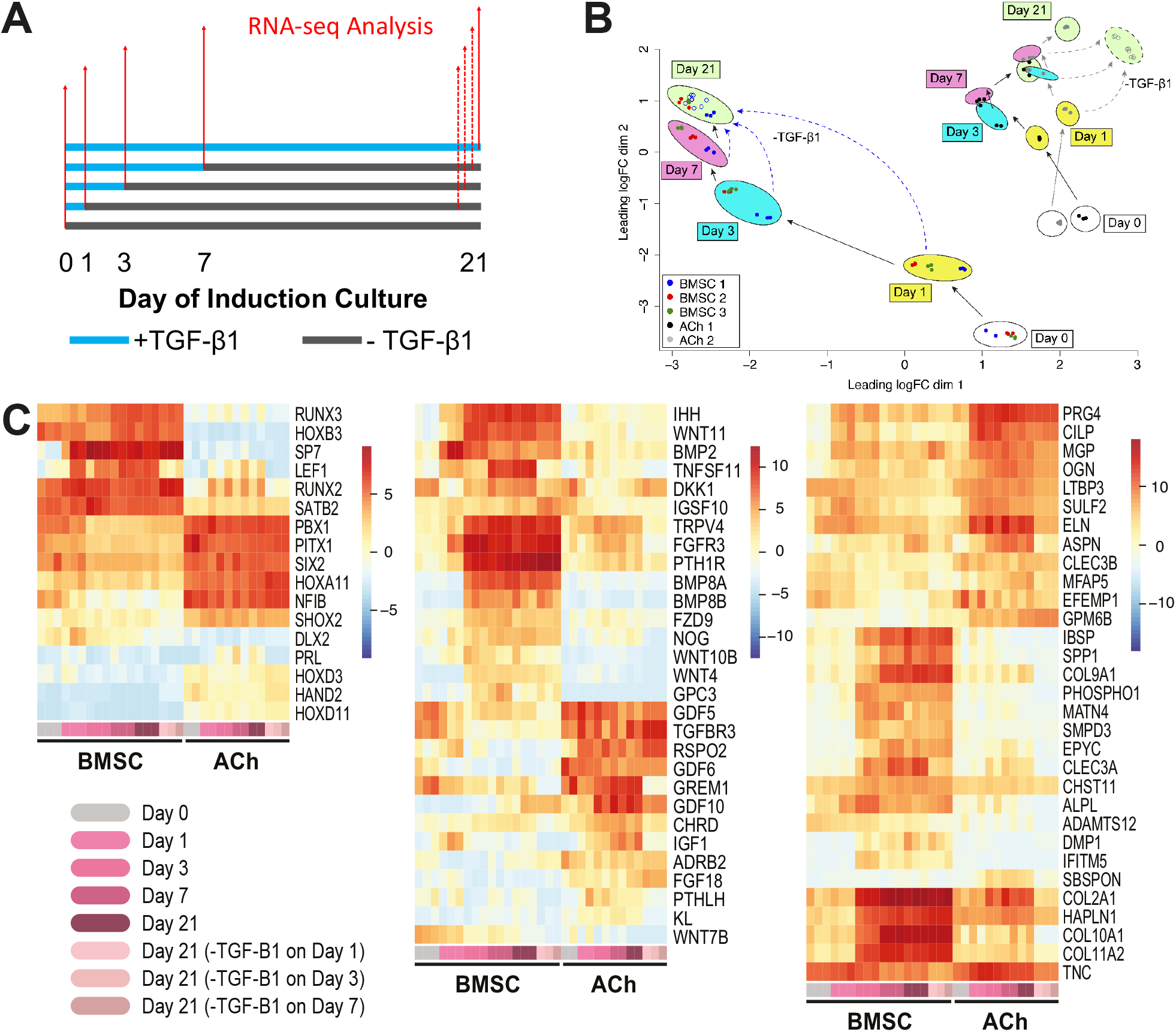
RNA-Sequencing analysis. (A) Whole transcriptome analysis was performed for BMSC and ACh cultures treated with TGF-β1 (blue lines) and harvested on day 0, 1, 3, 7 and 21 as indicated by the solid red arrows (BMSC donors 1, 2, and 3, and ACh donor 1 and 2). Additionally, for BMSC donor 1 and ACh donor 2, RNA-seq analysis was performed on day 21 following TGF-β1 withdrawal on day 1, 3, and 7, represented by dashed red arrows. (B) Multidimensional scaling (MDS) plot showing convergence of BMSC gene expression by day 21, despite withdrawal of exogenous TGF-β1 in some samples on day 1, 3, or 7, and dissimilarity to ACh samples treated in the same way. (C) Differentially expressed genes related to osteochondral transcription factors (left), soluble factors and receptor signalling (middle), and ECM molecules and ECM biosynthesis (right). Each timepoint in the heatmap is represented by a color shown on the bottom row of the heatmap, which corresponds to the color key on the bottom left of the figure. Each BMSC timepoint has 3 columns, representing 3 unique BMSC donors, and each ACh timepoint has 2 columns, representing 2 unique ACh donors; only one BMSC donor and one ACh donor was sequenced on day 21 with TGF-β1 withdrawal on day 1, 3, or 7, and are represented by the lighter pink shades; heatmap colors represent the average of three normalized logCPM values.

As shown in the multi-dimensional scaling (MDS) in **Fig 6B**, the three BMSC donors analyzed on a specific day after TGF-β1 treatment clustered well with each other at each time point. With incremental culture in TGF-β1-supplemented medium, BMSC gene expression shifted to the upper left corner of the plot for all BMSC donors. When 21-day cultures were replicated with BMSC donor 1 (open blue points), but with TGF-β1 eliminated from cultures at day 1, 3, 7, or 21, gene expression converged on a common signature identified in the green oval on the MDS plot. This clustering of day 21 cultures, regardless of the length of TGF-β1 exposure, confirmed that even with as little as 1 day of TGF-β1 exposure, BMSCs are fated to undergo a chondrogenic and hypertrophic differentiation program. Importantly, BMSCs and ACh did not converge in the MDS plot, showing dissimilarity in gene expression, despite being kept under identical culture conditions. While ACh that had been exposed to fewer days of TGF-β1 (open grey points) clustered closely together on day 21, they did not completely overlap with ACh that had been exposed to TGF-β1 for the full 21-day culture duration. This suggests that, unlike BMSCs, ACh might undergo a de-differentiation, or other, transcriptional program in response to the withdrawal of TGF-β1 signal.

At each of the timepoints that we performed RNA-seq, we analyzed the data for differentially expressed genes between BMSCs and ACh (see Supplementary Table 1). We aimed to produce a list of genes that could be useful for detecting early markers of BMSC hypertrophy or as targets to mitigate hypertrophy. We compiled a list of genes that were consistently differentially expressed (>2 log fold) between BMSC and ACh samples on days 3, 7, and 21, which resulted in a list of 947 genes. We assessed this list of genes against gene ontology terms related to cartilage development (GO:0051216) and ossification (GO:0001503) [16], reducing the list to 77 relevant genes. We further categorized this list into genes related to transcription factors, soluble factors and receptor signalling, and ECM molecules and ECM biosynthesis, and plotted the gene logCPM values for each sample to generate heatmaps (**Fig 6C**).

Genes for transcription factors that were expressed higher in BMSCs, compared with ACh, included *RUNX3*, *HOXB3*, *SP7*, *LEF1*, *RUNX2*, and *SATB2*. Of these, SP7 and LEF1 were specifically up-regulated in response to TGF-β1 exposure in BMSCs. However, unlike LEF1, SP7 remained highly expressed on day 21 even with the withdrawal of TGF-β1 on day 1, 3, or 7. Because SP7 was only highly expressed following TGF-β1 induction and maintained this level of expression even after TGF-β1 withdrawal, SP7 expression might be the most reliable early transcription factor to detect BMSC hypertrophic fate. A number of genes for transcription factors were significantly higher in ACh, compared with BMSCs, but these genes did not appear to change exclusively in response to TGF-β1 or its withdrawal. Many of the transcription factors that were expressed higher in ACh, including a number of *HOX* genes, *PBX1* [17], *PITX1* [18], *SIX2* [19], *NFIB* [20], and *SHOX2* [21], have been associated with patterning and early musculoskeletal progenitor cells.

Genes associated with soluble signalling factors and receptors that were differentially expressed between BMSCs and ACh were largely related to TGF-β superfamily signalling, the WNT signalling pathway, and a few others. *IHH*, *WNT11*, *WNT4* [22], and *BMP2* [23] are associated with cartilage hypertrophy, and in our study they were upregulated by a single day of TGF-β1 exposure in BMSCs, and their expression remained elevated following its withdrawal. *PTH1R* expression, which is associated with bone formation and resorption [24], was expressed significantly higher in BMSCs compared with ACh, in response to any duration of TGF-β1 exposure. ACh expressed higher levels of *PTHLH*, a known modulator of cartilage hypertrophy [25], than BMSC, in the presence of TGF-β1, but not following its withdrawal. Chondrocyte-associated factors from the TGF-β superfamily, *GDF5, GDF6* [26] and *TGFBR3* [27], were consistently expressed in ACh, but were under-expressed by TGF-β1 exposure in BMSCs. BMP antagonists *GREM1* and *CHRD* were expressed by both BMSCs and ACh prior to induction (Day 0) and in the presence of TGF-β1, with higher relative expression in ACh, and decreased in both cell types following TGF-β1 withdrawal. *FGF-18*, a mitigator of hypertrophy [28], was expressed higher in ACh than BMSCs at each timepoint and regardless of TGF-β1 exposure or following its withdrawal.

In both BMSCs and ACh, the expression of genes related to ECM molecules and ECM biosynthesis that were responsive to TGF-β1 were typically upregulated within either 1 day, or by 3 days, of TGF-β1 exposure, and generally remained upregulated following TGF-β1 withdrawal with some exceptions in ACh. In both BMSCs and ACh, *PRG4* and *CILP* were upregulated after 1 day of TGF-β1 exposure, with overall greater expression in ACh, even following TGF-β1 withdrawal. In BMSCs, but not ACh, genes associated with ECM mineralization were upregulated including *IBSP, SPP1*, and *PHOSPHO1*, but required more than 1 day of TGF-β1 exposure and remained highly expressed following its withdrawal. The upregulation of ECM molecule genes associated with cartilage and bone development including *COL2A1*, *HAPLN1*, *COL10A1*, and *COL11A2* required more than 1 day of TGF-β1 exposure in BMSCs, but remained highly expressed despite TGF-β1 withdrawal. While *COL2A1* and *ELN* expression in ACh was generally increased with prolonged TGF-β1 expression, TGF-β1 withdrawal significantly reduced the level of their expression. A low expression of the hypertrophic ECM molecule gene *COL10A1* was detected in ACh prior to induction and during TGF-β1 exposure, but this expression dropped dramatically following TGF-β1 withdrawal.

## Discussion

Limitations with ACh-based therapies for cartilage repair are driving investment into therapies that utilize BMSCs or other cell populations [29]. The most commonly evaluated alternative cell populations are bone marrow-derived cells (31%, including BMSCs) [29]. However, the progression of prospective therapies using bone marrow-derived cells to phase III trials is low (3.2% [29]), and no BMSC cartilage repair therapies have attained FDA approval. We reason that significant knowledge gaps in the understanding of BMSC chondrogenic differentiation play a major role in the failure of BMSCs to achieve cartilage repair expectations [1].

For over two decades, researchers have been using the pellet culture, in which several hundred-thousand BMSCs are pelleted in a tube containing medium supplemented with TGF-β1 ([6]; referred to as the *macro*-pellet in this manuscript) to study this differentiation process. However, this is a spatially and temporally heterogeneous model, which obfuscates the effects of the differentiation media.

Increasingly, the propensity of BMSCs to form hypertrophic tissue has been acknowledged as a barrier to successful clinical translation [1], and this has become an area of active research. Studies have reported successful obstruction of hypertrophy and formation of stable articular cartilage-like tissue by: (1) manipulation of WNT signalling [30, 31]; (2) silencing of BMP receptor signalling via a proprietary molecule [32]; (3) provision of different soluble factors on opposite sides of a cartilage-like disc to replicate an osteochondral interface [33]; or (4) use of novel scaffolds [34]. While promising, a recent article critically notes that these studies have yet to be replicated by independent groups [35]. A common feature of previous studies was the use of large tissue models and extended TGF-β exposure. For example, Narcisi *et al.*, used 2×10^5^ BMSC per *macro*-pellet and 5 weeks induction (10 ng/mL TGF-β1) [30], Yang *et al.*, 2.5×10^5^ BMSC per *macro*-pellet and 14-35 days induction (5 ng/mL TGF-β3) [31], Occhetta *et al.*, 2.5×10^5^ BMSC per *macro*-pellet and 14 days induction (10 ng/mL TGF-β3) [32], Ng *et al.*, multiplex layers of 2.5×10^5^ BMSC and 10 weeks induction (10 ng/mL TGF-β3) [33]. Physically large tissues, generated from hundreds of thousands of cells, suffer diffusion gradients resulting in spatial tissue heterogeneity. The desired stable high-quality cartilage-like tissue may have formed in regions of these *macro*-tissues, but likely tissue quality varied spatially.

Our group has been a proponent of using smaller diameter *micro*-pellets to generate more homogeneous tissues [4, 5], reasoning that a homogeneous readout is necessary for optimizing a complex bioprocess such as BMSC chondrogenesis. Here we studied BMSC differentiation in response to the canonical induction factor, TGF-β1, using a *micro*-pellet model (5×10^3^ BMSCs each) and a traditional *macro*-pellet model (2×10^5^ BMSCs each). We reasoned that in addition to the confounding heterogeneity of *macro*-pellet models, the use of cartilage-like matrix quantification as the readout for BMSC chondrogenesis can be misleading. Cartilage-like matrix deposition must necessarily lag behind the BMSC chondrogenic fate decision, introducing a significant delay in matrix readouts. As chondrocytes lie on a continuum between skeletal stem/progenitor cells, and hypertrophic chondrocytes, or bone-forming cells [1], a presumably temporal exposure of BMSCs to induction factors is likely to be a critical consideration in the optimisation of chondrogenesis protocols. To account for a lag in matrix deposition, we titrated the duration of TGF-β1 exposure (0, 1, 3, 7, 14, or 21 days), but maintained all cultures for 21 days to allow cell fate decisions to manifest and for matrix to accumulate

Physical tissue size provided an immediate indication that the kinetics of BMSC *micro*- and *macro*-pellets response to TGF-β1 exposure differed. As expected, extended TGF-β1 exposure resulted in incrementally larger BMSC *macro*-pellets. BMSC *macro*-pellet GAG and GAG/DNA quantities increased incrementally with extended TGF-β1 exposure. Unexpectedly, a single day of TGF-β1 exposure yielded BMSC *micro*-pellets that grew to a size nearly equivalent to *micro*-pellets exposed to TGF-β1 for 21 days. Biochemical characterization similarly indicated that brief exposure of BMSC *micro*-pellets to TGF-β1 was sufficient to yield GAG and GAG/DNA quantities similar to *micro*-pellets exposed to TGF-β1 for 21 days. These results suggested that either BMSCs responded differently to TGF-β1 in *micro*- and *macro*-pellet models, or that the model significantly influences the perceived kinetics.

Histological matrix characterisation of BMSC *micro*- or *macro*-pellets indicated that cells likely responded similarly to TGF-β1 exposure, but that the *micro*- or *macro*-pellet models significantly influenced perceived kinetics. Radially heterogenous GAG matrix deposition in *macro*-pellet histological sections suggested that BMSC differentiation occurred at different rates, depending on the location of the cells within the *macro*-pellet. Cartilage-like matrix formed first at the outer edge or in small localized regions of *macro*-pellets, converging incrementally throughout the *macro*-pellets if TGF-β1 exposure was extended. Heterogenous matrix accumulation observed in *macro*-pellets in our study is similar to previous reports [4, 5, 32], and perhaps rationalizes the common use of multi-week cultures with continuous TGF-β exposure. By contrast, when BMSCs were induced as *micro*-pellets, their response to TGF-β1 appeared more synchronized. A single day of TGF-β1 exposure resulted in *micro*-pellets having relatively uniform cartilage-like matrix distribution that was rich in GAG, and morphologically similar to *micro*-pellets exposed to TGF-β1 for 21 days. Similarly, lacunae structure appeared more mature in *micro*-pellets, compared with *macro*-pellets, exposed to a single day of TGF-β1. Cumulatively, these data indicate that reduced diffusion gradients in *micro*-pellets allow synchronized response to TGF-β1, revealing that a single day of TGF-β1 exposure is sufficient to trigger profound chondrogenic differentiation signaling cascades in BMSCs. By contrast, BMSC *macro*-pellets were spatially and temporally heterogenous, likely due to steep diffusion gradients. The relative response of ACh to TGF-β1 exposure in *micro*-vs. *macro*-pellet culture was not as profound as that of BMSCs. A single day of TGF-β1 exposure was insufficient to maximize the cartilage-like matrix output of 21 day ACh cultures. Both ACh *micro*- and *macro*-pellets benefited from extended TGF-β1 exposure, which increased both DNA and GAG content. Because ACh are a highly committed cell type, unlike multipotent BMSCs, it is possible that TGF-β1 serves more to stimulate matrix production in these cells rather than affect the cell phenotype.

We initially used qPCR to compare chondrogenic and hypertrophic gene expression in BMSC and ACh *micro*- and *macro*-pellets subjected to different durations of TGF-β1 exposure. Brief, 1-3 day exposure of BMSC *micro*-pellets to TGF-β1 was sufficient to upregulate both chondrogenic (*SOX9, COL2A1*, and *ACAN*), and hypertrophic (*COL10A1*) gene expression to levels seen in cultures exposed to TGF-β1 for the full 21 days of induction. By contrast, in ACh *micro*- and *macro*-pellets, chondrogenic genes were increased in response to incrementally greater TGF-β1 exposure, but this could only be observed with statistical significance for *ACAN* for one of the two ACh donors analyzed.

To better understand the transcriptional response to differential TGF-β1 exposure, we performed RNA-Seq on BMSCs and ACh *micro*-pellets. We analyzed data from cultures prior to induction, continuously exposed to TGF-β1 and harvested at day 1, 3, 7 or 21, as well as tissues cultured until day 21 after having been exposed to TGF-β1 for only 1, 3, or 7 days. This design provided insight into the temporal progression of the differentiation process, as well as opportunity to assess manifestation of programming triggered by different durations of TGF-β1 exposure. Each of the three unique BMSC donors generated a similar gene expression signature, progressively upregulating chondrogenic and hypertrophic genes with 1, 3, 7 or 21 days of TGF-β1 exposure. We assessed gene expression related to cartilage and bone development that was significantly differentially expressed more than 2 log fold between BMSCs and ACh following TGF-β1 exposure and categorized them by their function including those related to transcription factors, soluble signals and receptors, and ECM molecules and ECM biosynthesis.

SP7, or Osterix, is a transcription factor that drives hypertrophy [36] and is required for bone formation [37], and was found to be exclusively upregulated in BMSC in our study, with as little as 1 day of TGF-β1 exposure. One-day of TGF-β1 exposure resulted in a 9.7 log fold difference in *SP7* expression in BMSCs, compared with ACh cultures, and remained highly expressed following TGF-β1 withdrawal from BMSCs. *RUNX2*, the master regulator of osteoblast differentiation, was expressed in our BMSC cultures 6.4 log fold more than in ACh cultures prior to chondrogenic induction. This baseline expression of *RUNX2* does not increase significantly in BMSC following TGF-β1 exposure, suggesting that RUNX2 does not induce hypertrophy in BMSCs on its own, but likely requires TGF-β1 mediated upregulation of other transcription factors, such as SP7, for hypertrophic conversion. Previous studies have shown that knockdown of *Sp7* in mice results in diminished hypertrophy in bone progenitor cells *in vivo* [36]. While molecular inhibitors of *SP7* have not been reported, it would be of interest to determine whether regulation of *SP7* through knockdown in BMSCs could lead to high-quality, stable hyaline cartilage tissue that does not undergo hypertrophy.

We also identified the expression of soluble signalling factors associated with hypertrophy in BMSC belonging to the WNT and TGF-β1 superfamily signalling pathways, including BMP2 and WNT11. Inhibition of these factors in BMSC chondrogenic induction cultures has been previously reported with some success [32, 35]. Further improvement may be realized using combinations of these inhibitors in *micro*-pellet cultures as described here, at the time of pellet initiation, as hypertrophic fate commitment occurs with a single day of TGF-β1 exposure in BMSCs.

ECM molecules associated with BMSC hypertrophy and mineralization, such as *COL10A1* and *IBSP*, were not statistically differentially expressed between BMSC and ACh until more than 1-day of TGF-β1 exposure. ECM molecules and their biosynthesis are generally downstream products of transcriptional and receptor signalling machinery and, as such, it is not surprising that a relative delay in their gene expression might be observed. Expression of ECM associated genes can serve as good markers for assessing cartilage hypertrophy and tissue quality, but upstream targets are more likely, and feasible, to successfully mitigate hypertrophy.

Comparative analysis of RNA-Seq data from BMSC and ACh *micro*-pellets provides a number of useful insights into the pathways that differ between these cell populations, identifies potential target pathways to obstruct BMSC hypertrophy, and genes that could be used in reporter assays to facilitate development of chondrogenic media, cell processing, and scaffolding. Our results suggest that BMSC chondrogenic and hypertrophic differentiation is induced following a brief 1-day exposure to TGF-β1. Future efforts to generate stable cartilage-like tissue should focus on manipulation of the culture conditions over the first few hours of culture induction, including early efforts to obstruct hypertrophy. It is likely unrealistic to expect to identify and characterize these early events using a heterogenous readout, like a *macro*-pellet or macroscopic tissue model(s). While it may be necessary to develop macroscopic scaffold or gel delivery solutions for cartilage defect repair, addressing basic biological questions will require models not confounded by spatial heterogeneity. More sensitive or responsive models, such as the *micro*-pellet, will enable bioprocess approaches [38] to be exploited, enabling systematic progression towards optimal culture conditions. Generating high-quality hyaline cartilage is likely to require multiple factors in parallel or in sequence. Previous studies examined the influence of sequential growth factor stimulation at different days in *macro*-pellet cultures [11, 39, 40], likely overshooting the period where BMSCs are most sensitive to instruction. Using a *micro*-pellet model, or other homogeneous model, focused on guiding cell fate over the first hours or days of culture is more likely to reveal the growth factor combinations required to generate high-quality stable cartilage tissue.

## Conclusion

While BMSCs may have significant unrealized potential in cartilage tissue engineering, current chondrogenic differentiation protocols yield sub-optimal cartilage-like tissue with a hypertrophic propensity [1]. The majority of BMSC chondrogenesis studies characterise the influence of growth factors alone or in combination over multiple days or multiple weeks of culture. Using a *micro*-pellet model, we discovered that BMSCs respond to a single day of the canonical signalling molecule, TGF-β1, engaging both chondrogenic and hypertrophic differentiation pathways. This discovery is significant as it identifies that minimal TGF-β1 is required to engage this differentiation process, and that culture optimisation should focus on the first few hours or days of induction. Using more homogeneous small diameter tissue culture models, it may be possible to overcome current limitations in BMSC-mediated cartilage defect repair.

## Materials and Methods

### Human BMSC isolation and expansion

BMSC cultures were established using 20 mL of heparinized bone marrow aspirate (BMA) collected from the iliac crests of consenting normal adult human donors at the Mater Hospital, Brisbane, Australia. Ethics approval for aspirate collection was granted by the Mater Health Services Human Research Ethics Committee and the Queensland University of Technology Human Ethics Committee (Ethics number: 1000000938). BMSCs were collected as previously described [5] with the exception of donor 3, which was not enriched for mononuclear cells prior to plastic attachment overnight. BMSC donor details were as follows: 24-year-old male (BMSC 1), 44-year-old male (BMSC 2), 21-year-old female (BMSC 3), and 43-year-old male (BMSC 4). BMSC expansion medium contained low glucose DMEM with GlutaMAX and pyruvate, 10% fetal bovine serum (FBS), 100 U/ ml penicillin/streptomycin (PenStrep), all from Thermo Fisher, 10 ng/mL fibroblast growth factor-1 (FGF-1; Peprotech), and 5 μg/mL porcine heparin sodium salt (Sigma). Cells were distributed into five T175 culture flasks with 35 mL of expansion medium per flask and were allowed to attach to the plastic surface overnight in a 20% O_2_, 5% CO_2_ and 37 °C incubator. The medium was replaced after 24 hours to remove loose cells and cell expansion was continued in a reduced oxygen atmosphere, 2% O_2_, 5% CO_2_ and 37 °C incubator. At 80% confluence, cells were passaged using 0.25% trypsin/EDTA (Thermo Fisher) and fresh flasks were re-seeded at ~1,500 cells per cm^2^. All BMSC donor cells had undergone cryopreservation in 90% FBS and 10% DMSO, thawed, and induced at passage 3.

### Human articular chondrocyte expansion

Normal human articular chondrocytes (ACh) from the knee were purchased from Lonza. Based on the manufacturer’s information sheet the cells were cryopreserved at passage 2, and donor information was as follows: 34-year-old male (Lot#: BF3339; ACh 1 in this study) and 50-year-old male (Lot#: BF3307; ACh 2). After thawing, these cells were grown as described for BMSCs above, in a 2% O_2_, 5% CO_2_ and 37 °C incubator and induced at passage 6.

### Chondrogenic-induction cultures

Chondrogenic induction media consisted of high glucose DMEM supplemented with GlutaMAX and pyruvate, 1× insulin-transferrin-selenium-ethanolamine (ITS-X), PenStrep, all from Thermo Fisher, 200 μM ascorbic acid 2-phosphate (Sigma), 40 μg/mL l-proline (Sigma), and 10 ng/mL TGF-β1 (Peprotech) where specified. The cells in the “Day 0” TGF-B1 removal condition were never supplemented with TGF-β1. The cells were force-aggregated in the well plates by centrifugation at 500 × *g* for 3 minutes. Full medium exchanges were performed every 2 days or on specified day of TGF-β1 removal. For TGF-β1 removal, the cell pellets were rinsed once with PBS, then fresh induction medium without TGF-β1 was added.

For *macro*-pellet cultures, cells were seeded in 96-well plates containing deep v-bottom wells (Sigma, Cat#: 3960), such that each well or aggregate contained 2×10^5^ cells per well in 1 mL of culture medium. The construction of microwell-mesh plates, including necessary materials, plate sterilisation and characterization is described in detail in [5]. These plates contain an array of polydimethylsiloxane (PDMS) pyramidal microwells (2 × 2 × 0.8 mm), covered with a nylon (6/6) mesh (36 μm pore opening). The PDMS microwells enable the formation of many cellular *micro*-pellets, simultaneously, while the nylon mesh prevents displacement of the *micro*-pellets from microwells over long-term culture. An animation demonstrating how the Microwell-mesh platform functions to efficiently manufacture hundreds of *micro*-pellets is available here [41]. In *micro*-pellet cultures, cells were seeded at 1.25×10^6^ per well of a 6-well plate. At this seeding density, ~250 *micro*-pellets are formed in each well, with each *micro*-pellet containing ~5,000 BMSC. Each well contained 4 mL of total culture medium.

### GAG and DNA quantification

GAG and DNA was analyzed as previously described [5]. Briefly, tissues were papain (Sigma) digested overnight at 60 °C. GAG was quantified using the 1,9-dimethymethylene blue (Sigma) assay, using chondroitin sulfate from shark cartilage (Sigma) to generate a standard curve. DNA was quantified using the PicoGreen assay kit (Thermo Fisher). Four replicate samples were analysed for each unique cell donor and the means were compared to respective Day 21 +TGF-β1 control cultures.

### RNA Isolation

Samples were crushed in RLT buffer (Qiagen) containing β-mercaptoethanol (Sigma) and RNA was isolated using the RNeasy Mini Kit (Qiagen) with on-column DNase I (Qiagen) digestion, as per manufacturer’s instructions. RNA concentrations were determined using a NanoDrop 1000 (Thermo Fisher).

### Quantitative-PCR (qPCR) Analysis

cDNA was synthesized from total RNA using SuperScript III First-Strand Synthesis System for RT-PCR (Thermo Fisher), as per manufacturer’s instructions. qPCR reactions were prepared using SYBR Green PCR Master Mix (Applied Biosystems). Three technical replicates were loaded for each of the 4 biological replicates per donor in 384 well plates and analysed on a Viia7 Real Time PCR System (Applied Biosystems). The forward and reverse primer sequence for target genes, as well as the qPCR run parameters have been previously published [5]. Target gene expression was normalized to GAPDH expression and calculated using the ΔCt method. Statistically significant differences in relative gene expression levels for each donor were determined using one-way ANOVA in GraphPad Prism 7.0.

### Subcutaneous implantation of cartilage defect model

NOD/LtSz-scid IL2R gamma null (NSG) mice were purchased from the Jackson Laboratory and bred in the Animal Facility at the Translational Research Institute (TRI) in Brisbane. The University of Queensland (UQ) and the Queensland University of Technology (QUT) Animal Ethics Committees authorized the animal procedures described here. All animal procedures were carried out in accordance with the approved guidelines (Ethics numbers: AEMAR53777 and AEMAR53765). Artificial cartilage defect models were prepared from bovine tissue as previously described [5]. Briefly, plugs containing cartilage and bone were drilled out from bovine knees using a 10 mm coring bit. Full thickness cartilage defects were drilled out of the plug using a 3.5 mm drill bit. The defects were washed of debris and sterilized in 70% ethanol for 24 hours and washed 3× in PBS over another 24 hours. The PBS was discarded, and the defects were kept frozen at −20 °C until use. The defects were filled with 21-day cultured *micro*-pellets that were grown in the presence of TGF-β1 for 21 days, or had TGF-β1 withdrawn at earlier timepoints, and sealed with fibrin glue (one drop of fibrinogen, followed by one drop of thrombin) (Tisseel, Baxter). Each defect was implanted subcutaneously in a pocket made on the back of an anaesthetized NSG mouse and the skin stapled to close the wound. These tissues were permitted to incubate in NSG mice for 8 weeks, at which point the animals were euthanized and the tissues recovered for analysis.

### microCT Analysis

Tissues excised from mice were fixed in 4% paraformaldehyde (PFA) for 24 hours and scanned in plastic tubes containing 70% alcohol and styrofoam to reduce movement. MicroCT analysis was performed to detect hard tissue formation using an Inveon PET-CT Scanner (Siemens; 27.6 μm pixel, 60 kV, 350 μA, 2.5 s exposure) or the SkyScan 1272 (Bruker; 17-22 μm pixel, 50 kV, 200 μA, 0.25 mm Al filter, 425 ms exposure). Reconstruction and imaging was performed using Inveon Software (Siemens) or NRecon Software and DataViewer Sofware (Skyscan Bruker) for related machines.

### Histology

Tissues induced for 21 days were fixed in 4% PFA for 30 minutes, washed and frozen in Tissue-Tek OCT compound (Sakura Finetek). Samples were cryosectioned at 7 μm and collected onto polylysine coated slides (Thermo Fisher). After microCT analysis, tissues excised from mice were decalcified in EDTA solution until soft, dehydrated, paraffin embedded and sectioned at 5 μm. Sections were stained with Toluidine blue (Sigma) for glycosaminoglycan (GAG).

### RNA-Seq

Total RNA was collected from BMSC and ACh cultures prepared in microwell-mesh plates on day 0, 1, 3, 7 and 21, with or without TGF-β1 washout, as indicated. “Day 0” cells represent expanded cells that were not induced. RNA integrity was confirmed using the Agilent 2100 Bioanalyser (Agilent Technologies). Next-generation sequencing and primary bioinformatics was performed by the Australian Genome Research Facility (AGRF, Melbourne) using the Illumina HiSeq 2500 platform (100 bp single end run). AGRF analysis involved demultiplexing, quality control, and then data was processed through RNA-seq expression analysis workflow, which included alignment, transcript assembly, quantification, normalisation, and differential gene expression analysis (Bioconductor R package edgeR [42]). Significance of differentially expressed genes between ACh and BMSC cultures was considered at an adjusted *p-value* (FDR) of <0.05. Heatmaps were generated using the heatmap.2 function in the R gplots package [43].

### Statistical Analysis

The quantification of GAG/DNA and qPCR are presented as mean ± SD (n = 4 for each cell donor). Statistical analysis was performed using ANOVA in GraphPad Prism version 7. P-values of 0.05 or less were considered statistically significant.

## Data Availability

Data supporting the conclusions of this paper are available from the corresponding author upon request.

## Acknowledgments

The Translational Research Institute (TRI) is supported by Therapeutic Innovation Australia (TIA). TIA is supported by the Australian Government through the National Collaborative Research Infrastructure Strategy (NCRIS) program. KF and MRD thank the TRI Biological Resource Facility for help with animal studies, the TRI Preclinical Imaging Facility for help with microCT analysis, the TRI Histology Facility for help with tissue processing and sectioning, the Mater Hospital for BMA collection, and the Australian Genome Research Facility (AGRF) for RNA-seq and bioinformatics analysis. KF and MRD gratefully acknowledge project support from the National Health and Medicine Research Council (NHMRC) of Australia (Project Grant APP1083857) and NHMRC Fellowship support of MRD (APP1130013). PGR is supported by the Division of Intramural Research (DIR) of the National Institute of Dental and Craniofacial Research (NIDCR), a part of the Intramural Research Program (IRP) of the National Institutes of Health (NIH), Department of Health and Human Services (DHHS) (1 ZIA DE000380 35). The authors would like to thank Ms Ena Music and abpLearning for generously producing the Microwell-mesh animation.

## References

1. Somoza, R.A., J.F. Welter, D. Correa, and A.I. Caplan, Chondrogenic differentiation of mesenchymal stem cells: challenges and unfulfilled expectations. Tissue Eng Part B Rev, 2014. 20(6): p. 596–608.

2. Sacchetti, B., A. Funari, S. Michienzi, S. Di Cesare, S. Piersanti, I. Saggio, E. Tagliafico, S. Ferrari, P.G. Robey, M. Riminucci, and P. Bianco, Self-renewing osteoprogenitors in bone marrow sinusoids can organize a hematopoietic microenvironment. Cell, 2007. 131(2): p. 324–36.

3. Hunziker, E.B., T.M. Quinn, and H.J. Hauselmann, Quantitative structural organization of normal adult human articular cartilage. Osteoarthritis Cartilage, 2002. 10(7): p. 564–72.

4. Markway, B.D., G.K. Tan, G. Brooke, J.E. Hudson, J.J. Cooper-White, and M.R. Doran, Enhanced chondrogenic differentiation of human bone marrow-derived mesenchymal stem cells in low oxygen environment micropellet cultures. Cell Transplant, 2010. 19(1): p. 29–42.

5. Futrega, K., J.S. Palmer, M. Kinney, W.B. Lott, M.D. Ungrin, P.W. Zandstra, and M.R. Doran, The microwell-mesh: A novel device and protocol for the high throughput manufacturing of cartilage microtissues. Biomaterials, 2015. 62: p. 1–12.

6. Johnstone, B., T.M. Hering, A.I. Caplan, V.M. Goldberg, and J.U. Yoo, In vitro chondrogenesis of bone marrow-derived mesenchymal progenitor cells. Exp Cell Res, 1998. 238(1): p. 265–72.

7. Freyria, A.M. and F. Mallein-Gerin, Chondrocytes or adult stem cells for cartilage repair: the indisputable role of growth factors. Injury, 2012. 43(3): p. 259–65.

8. Goldberg, A., K. Mitchell, J. Soans, L. Kim, and R. Zaidi, The use of mesenchymal stem cells for cartilage repair and regeneration: a systematic review. J Orthop Surg Res, 2017. 12(1): p. 39.

9. Jakobsen, R.B., E. Ostrup, X. Zhang, T.S. Mikkelsen, and J.E. Brinchmann, Analysis of the effects of five factors relevant to in vitro chondrogenesis of human mesenchymal stem cells using factorial design and high throughput mRNA-profiling. PLoS One, 2014. 9(5): p. e96615.

10. Puetzer, J.L., J.N. Petitte, and E.G. Loboa, Comparative review of growth factors for induction of three-dimensional in vitro chondrogenesis in human mesenchymal stem cells isolated from bone marrow and adipose tissue. Tissue Eng Part B Rev, 2010. 16(4): p. 435–44.

11. Handorf, A.M. and W.J. Li, Induction of mesenchymal stem cell chondrogenesis through sequential administration of growth factors within specific temporal windows. J Cell Physiol, 2014. 229(2): p. 162–71.

12. Vinatier, C., D. Mrugala, C. Jorgensen, J. Guicheux, and D. Noel, Cartilage engineering: a crucial combination of cells, biomaterials and biofactors. Trends Biotechnol, 2009. 27(5): p. 307–14.

13. McMurtrey, R.J., Analytic Models of Oxygen and Nutrient Diffusion, Metabolism Dynamics, and Architecture Optimization in Three-Dimensional Tissue Constructs with Applications and Insights in Cerebral Organoids. Tissue Eng Part C Methods, 2016. 22(3): p. 221–49.

14. Akkerman, N. and L.H. Defize, Dawn of the organoid era: 3D tissue and organ cultures revolutionize the study of development, disease, and regeneration. Bioessays, 2017. 39(4).

15. Babur, B.K., M. Kabiri, T.J. Klein, W.B. Lott, and M.R. Doran, The rapid manufacture of uniform composite multicellular-biomaterial micropellets, their assembly into macroscopic organized tissues, and potential applications in cartilage tissue engineering. PLoS One, 2015. 10(5): p. e0122250.

16. Jax. Mouse Genome Database (MGD), Gene Ontology (GO). (Accessed on 20 Oct 2019); Available from: http://www.informatics.jax.org/vocab/gene_ontology/.

17. Selleri, L., M.J. Depew, Y. Jacobs, S.K. Chanda, K.Y. Tsang, K.S. Cheah, J.L. Rubenstein, S. O’Gorman, and M.L. Cleary, Requirement for Pbx1 in skeletal patterning and programming chondrocyte proliferation and differentiation. Development, 2001. 128(18): p. 3543–57.

18. DeLaurier, A., R. Schweitzer, and M. Logan, Pitx1 determines the morphology of muscle, tendon, and bones of the hindlimb. Dev Biol, 2006. 299(1): p. 22–34.

19. He, G., S. Tavella, K.P. Hanley, M. Self, G. Oliver, R. Grifone, N. Hanley, C. Ward, and N. Bobola, Inactivation of Six2 in mouse identifies a novel genetic mechanism controlling development and growth of the cranial base. Dev Biol, 2010. 344(2): p. 720–30.

20. Harris, L., L.A. Genovesi, R.M. Gronostajski, B.J. Wainwright, and M. Piper, Nuclear factor one transcription factors: Divergent functions in developmental versus adult stem cell populations. Dev Dyn, 2015. 244(3): p. 227–38.

21. Bobick, B.E. and J. Cobb, Shox2 regulates progression through chondrogenesis in the mouse proximal limb. J Cell Sci, 2012. 125(Pt 24): p. 6071–83.

22. Hartmann, C. and C.J. Tabin, Dual roles of Wnt signaling during chondrogenesis in the chicken limb. Development, 2000. 127(14): p. 3141–59.

23. Church, V., T. Nohno, C. Linker, C. Marcelle, and P. Francis-West, Wnt regulation of chondrocyte differentiation. J Cell Sci, 2002. 115(Pt 24): p. 4809–18.

24. Datta, N.S. and A.B. Abou-Samra, PTH and PTHrP signaling in osteoblasts. Cell Signal, 2009. 21(8): p. 1245–54.

25. Minina, E., C. Kreschel, M.C. Naski, D.M. Ornitz, and A. Vortkamp, Interaction of FGF, Ihh/Pthlh, and BMP signaling integrates chondrocyte proliferation and hypertrophic differentiation. Dev Cell, 2002. 3(3): p. 439–49.

26. Chen, H., T.D. Capellini, M. Schoor, D.P. Mortlock, A.H. Reddi, and D.M. Kingsley, Heads, Shoulders, Elbows, Knees, and Toes: Modular Gdf5 Enhancers Control Different Joints in the Vertebrate Skeleton. PLoS Genet, 2016. 12(11): p. e1006454.

27. Wang, W., D. Rigueur, and K.M. Lyons, TGFbeta signaling in cartilage development and maintenance. Birth Defects Res C Embryo Today, 2014. 102(1): p. 37–51.

28. Liu, Z., J. Xu, J.S. Colvin, and D.M. Ornitz, Coordination of chondrogenesis and osteogenesis by fibroblast growth factor 18. Genes Dev, 2002. 16(7): p. 859–69.

29. Negoro, T., Y. Takagaki, H. Okura, and A. Matsuyama, Trends in clinical trials for articular cartilage repair by cell therapy. NPJ Regen Med, 2018. 3: p. 17.

30. Narcisi, R., M.A. Cleary, P.A. Brama, M.J. Hoogduijn, N. Tuysuz, D. ten Berge, and G.J. van Osch, Long-term expansion, enhanced chondrogenic potential, and suppression of endochondral ossification of adult human MSCs via WNT signaling modulation. Stem Cell Reports, 2015. 4(3): p. 459–72.

31. Yang, Z., Y. Zou, X.M. Guo, H.S. Tan, V. Denslin, C.H. Yeow, X.F. Ren, T.M. Liu, J.H. Hui, and E.H. Lee, Temporal activation of beta-catenin signaling in the chondrogenic process of mesenchymal stem cells affects the phenotype of the cartilage generated. Stem Cells Dev, 2012. 21(11): p. 1966–76.

32. Occhetta, P., S. Pigeot, M. Rasponi, B. Dasen, A. Mehrkens, T. Ullrich, I. Kramer, S. Guth-Gundel, A. Barbero, and I. Martin, Developmentally inspired programming of adult human mesenchymal stromal cells toward stable chondrogenesis. Proc Natl Acad Sci U S A, 2018. 115(18): p. 4625–4630.

33. Ng, J.J., Y. Wei, B. Zhou, J. Bernhard, S. Robinson, A. Burapachaisri, X.E. Guo, and G. Vunjak-Novakovic, Recapitulation of physiological spatiotemporal signals promotes in vitro formation of phenotypically stable human articular cartilage. Proc Natl Acad Sci U S A, 2017. 114(10): p. 2556–2561.

34. Kuznetsov, S.A., A. Hailu-Lazmi, N. Cherman, L.F. de Castro, P.G. Robey, and R. Gorodetsky, In Vivo Formation of Stable Hyaline Cartilage by Naive Human Bone Marrow Stromal Cells with Modified Fibrin Microbeads. Stem Cells Transl Med, 2019. 8(6): p. 586–592.

35. Diederichs, S., V. Tonnier, M. Marz, S.I. Dreher, A. Geisbusch, and W. Richter, Regulation of WNT5A and WNT11 during MSC in vitro chondrogenesis: WNT inhibition lowers BMP and hedgehog activity, and reduces hypertrophy. Cell Mol Life Sci, 2019. 76(19): p. 3875–3889.

36. Kaback, L.A., Y. Soung do, A. Naik, N. Smith, E.M. Schwarz, R.J. O’Keefe, and H. Drissi, Osterix/Sp7 regulates mesenchymal stem cell mediated endochondral ossification. J Cell Physiol, 2008. 214(1): p. 173–82.

37. Rashid, H., C. Ma, H. Chen, H. Wang, M.Q. Hassan, K. Sinha, B. de Crombrugghe, and A. Javed, Sp7 and Runx2 molecular complex synergistically regulate expression of target genes. Connect Tissue Res, 2014. 55 Suppl 1: p. 83–7.

38. Lipsitz, Y.Y., P. Bedford, A.H. Davies, N.E. Timmins, and P.W. Zandstra, Achieving Efficient Manufacturing and Quality Assurance through Synthetic Cell Therapy Design. Cell Stem Cell, 2017. 20(1): p. 13–17.

39. Indrawattana, N., G. Chen, M. Tadokoro, L.H. Shann, H. Ohgushi, T. Tateishi, J. Tanaka, and A. Bunyaratvej, Growth factor combination for chondrogenic induction from human mesenchymal stem cell. Biochem Biophys Res Commun, 2004. 320(3): p. 914–9.

40. Correa, D., R.A. Somoza, P. Lin, S. Greenberg, E. Rom, L. Duesler, J.F. Welter, A. Yayon, and A.I. Caplan, Sequential exposure to fibroblast growth factors (FGF) 2, 9 and 18 enhances hMSC chondrogenic differentiation. Osteoarthritis Cartilage, 2015. 23(3): p. 443–53.

41. Futrega, K. Microwell-mesh animation. 2019 [cited 2019 November 29]; Available from: https://youtu.be/mhp8k_bGtCw

42. Robinson, M.D., D.J. McCarthy, and G.K. Smyth, edgeR: a Bioconductor package for differential expression analysis of digital gene expression data. Bioinformatics, 2010. 26(1): p. 139–40.

43. Warnes, G.R.B., B.; Bonebakker, L.; Gentleman, R.; Huber, W.; Liaw, A.; Lumley, T.; Maechler, M.; Magnusson, A.; Moeller, S.; Schwartz, M.; Venables, B. gplots: Various R Programming Tools for Plotting Data. Version 3.0.1.1. 2015 (Accessed on 20 Oct 2019); Available from: https://cran.r-project.org/package=gplots.

